## Supplementary figures and images for "Super-enhancer-driven ZFP36L1 promotes PD-L1 expression in infiltrative gastric cancer"

### Supplementary Figure 1

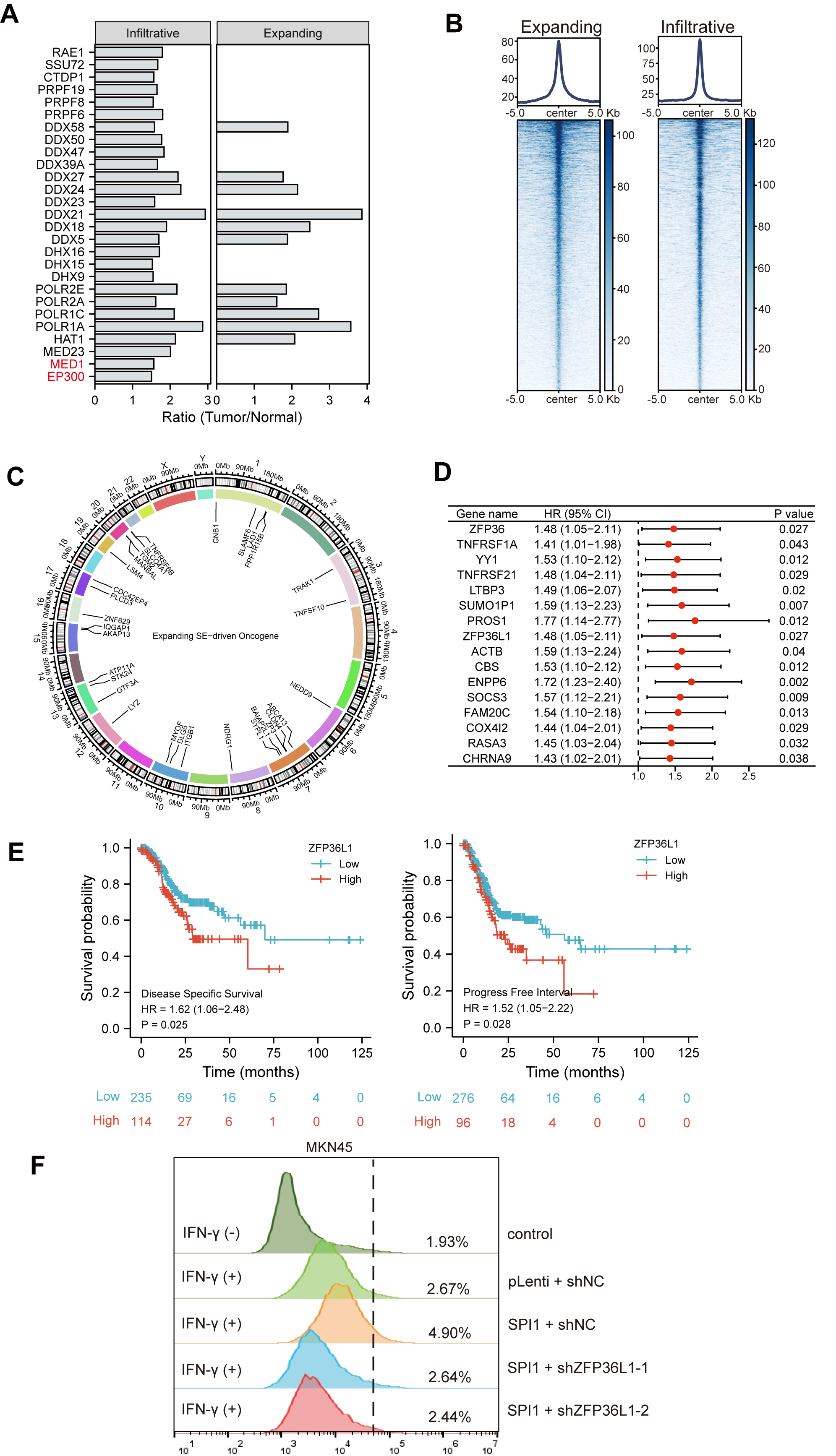
