## Supplementary Files for "Super-enhancer-driven ZFP36L1 promotes PD-L1 expression in infiltrative gastric cancer"

**1.Table 1**

| GENE | 5’ to 3’ |
| --- | --- |
| GAPDH | Forward: ACAACTTTGGTATCGTGGAAGG  Reverse: GCCATCACGCCACAGTTTC |
| ZFP36L1 | Forward: ACCACCACCCTCGTGTCTG  Reverse: TGCCCACTGCCTTTCTGT |
| CD274 | Forward: TGGCATTTGCTGAACGCATTT  Reverse: TGCAGCCAGGTCTAATTGTTTT |
| HDAC3 | Forward: CCTGGCATTGACCCATAGCC  Reverse: CTCTTGGTGAAGCCTTGCATA |
| SPI1 | Forward: ATGGAAGGGTTTCCCCTCGT  Reverse: CTGGAGCTCCGTGAAGTTGT |
| 18S rRNA | Forward: GTCTGTGATGCCCTTAGATG  Reverse: AGCTTATGACCCGCACTTAC |
| Mouse ZFP36L1 | Forward: CACCCCAAGTACAAGACGGA  Reverse: GCTAGGAGCAAAGAGGCTCG |
| Mouse CD274 | Forward: GCTCCAAAGGACTTGTACGTG  Reverse: TGATCTGAAGGGCAGCATTTC |
| ChIP-ZFP36L1-E1A | Forward: AAGTGCCAGTTTTCTTCCTTG  Reverse: CACCAGTCCCTGCCAGTC |
| ChIP-ZFP36L1-E1B | Forward: CTAGCAAGGCCCTGGTATG  Reverse: CTGTCCACATGGCAACCCT |
| ChIP-ZFP36L1-E1C | Forward: TTATACAACGTGGTGCTGGTG  Reverse: GTGTCAGTGCCTCCTCATT |
| ChIP-ZFP36L1-E1D | Forward: GGAGGCACTGACACGGACA  Reverse: GAATTCAAGTGGGGATTAGG |
| ChIP-CD274-P1 | Forward: GCTGGGCCCAAACCCTATT  Reverse: TTTGGCAGGAGCATG GAGTT |
| ChIP-CD274-P2 | Forward: ATGGGTCTGCTGCTGACTTT  Reverse: GGCGTCCCCCTTTCTGATAA |
| ChIP-CD274-P3 | Forward: ACTGAAAGCTTCCGCCGATT  Reverse: CCCAAGGCAGCAAAT CCAGT |

2. **Plasmid**

pLenti-CMV-ZFP36L1-GFP-Puro

pcDNA3.1-ZFP36L1-Flag

pPLK-GFP-Puro-ZFP36L1 shRNA-1: GTAACAAGATGCTCAACTATA

pPLK-GFP-Puro-ZFP36L1 shRNA-2: CCTCCAGCATAGCTTTAGCTT

pcDNA3.1-mutZFP36L1-Flag (C153R-C173R)

pLVX-CMV-ZFP36L1(mouse)-3×Flag-Puro

pLKO.1-U6-ZFP36L1 shRNA-1(mouse)-EF1a-copGFP-T2A-Puro: GCTTTCGAG

ACCGCTCTTTCTC

pLKO.1-U6-ZFP36L1 shRNA-2(mouse)-EF1a-copGFP-T2A-Puro: GCTGCCACTT

CATTCATAACGC

pLenti-CMV-SPI1-GFP-Puro——ID：6688；NM_001080547.2

pLenti-CMV-ELF1-GFP-Puro——ID：1997；NM_172373.4

pLenti-CMV-E2F1-GFP-Puro——ID：1869；NM_005225.3

pcDNA3.1-BRD4-3×Flag

pGEX-4T-2-GST-SPI1

pET-32a-His-BRD4

pLVX-Puro-Flag-HDAC3

pGL4-Luci-E1 (Full length)——chr14:68806839-68807740

pGL4-Luci-E1A——chr14:68806839-68807000

pGL4-Luci-E1B——chr14:68807000-68807300

pGL4-Luci-E1C (Wild)——chr14:68807300-68807500

pGL4-Luci-E1C (Deletion)——chr14:68807300-68807469 + 68807483-68807500, deletion: GAAGAGGGAAGGCAG

pGL4-Luci-E1D——chr14:68807500-68807740

pGL4-Luci-CD274 promoter

pmirGLO-HDAC3-3’UTR (Wild)——“ATTTA” motif

pmirGLO-HDAC3-3’UTR (Mutant)——“ACCCA” mutant motif

3. **Materials and Methods**
   3.1 **Statistical methods**

If the variable is numerical and the sample size is ≤5000, a normality test will be conducted. If the data follow a normal distribution, the mean ± standard deviation of the corresponding variable will be calculated; otherwise, the median (upper and lower quartiles) of the corresponding variable will be reported. For numerical variables that satisfy the normal distribution and pass the variance chi-square test, two-group comparisons will be performed using the T-test, and three-group comparisons will use One-way ANOVA. If the data satisfy the normal distribution but fail the variance chi-square test, two-group comparisons will be conducted using Welch's t-test, and three-group comparisons will use Welch's one-way ANOVA. If the data do not satisfy the normal distribution assumption, two-group comparisons will use Welch's one-way ANOVA. For normally distributed data, two-group comparisons will use the Wilcoxon test, and three-group comparisons will use the Kruskal-Wallis test.

If the variables are categorical and the data satisfy the condition of theoretical frequency > 5 with a total sample size ≥ 40, group comparisons will use the chi-square test. When the data meet the condition of 1 ≤ theoretical frequency ≤ 5 and the total sample size ≥ 40, group comparisons will use the corrected chi-square test (Yates' correction). If the data do not meet the conditions of theoretical frequency <1 or total sample size <40, group comparisons will utilize Fisher's exact test.

3.2 **Hematoxylin & eosin (HE) and immunohistochemical (IHC) staining**

3.3.1 Tissue Preparation:

Fresh tissue specimens are immersed in formalin fixative solution and incubated overnight at 4°C on a shaker. Tissues are dehydrated using an automatic dehydration instrument, embedded in paraffin using a paraffin embedding machine, and stored as tissue blocks at -20°C. Tissue sections of 4 μm thickness are prepared using a microtome, followed by flattening, lifting, and drying in a dark place.

3.3.2 Deparaffinization:

Slides are baked at 65°C in a constant temperature chamber for 6-24 hours.

Before staining, slides are deparaffinized to water. This involves sequential immersion in decreasing concentrations of ethanol: xylene substitute for 10 minutes, absolute ethanol for 2 minutes, 95% ethanol for 2 minutes, 75% ethanol for 2 minutes, and finally pure water, with each step repeated twice. For Hematoxylin-Eosin (HE) staining, slides are directly stained with hematoxylin for 3 minutes, followed by 10 seconds of decolorization in hydrochloric acid ethanol and rinsing in pure water for 5 minutes. Then, slides are stained with eosin for 30 seconds before proceeding to step 5 for mounting.

3.3.3 Antigen Retrieval:

Antigen retrieval is performed using high-pressure steam. Slides are placed in a metal dish filled with sodium citrate buffer solution and completely submerged. The dish is then placed in a preheated boiling pressure cooker and sealed. The heating plate is set to maximum power. Once the pressure cooker reaches maximum pressure and the valve releases steam for 2 minutes, the heating plate is turned off. After waiting for 30 minutes, the lid is opened, and the slides are cooled at room temperature for 2 hours.

3.3.4 Immunostaining:

Immunohistochemical experiments are conducted using an immunostaining kit. The specific steps are as follows: Wash the fixed sections with PBS solution for 3 minutes × 2 times, then rinse with PBS for the third time. Apply endogenous peroxidase blocker to the defined tissue area on the slide and incubate at room temperature for 10 minutes. Wash the slides with PBS for 3 minutes × 3 times, remove PBS, apply nonspecific staining blocker, and incubate at room temperature for 10 minutes. Apply primary antibody (diluted at 1:500) and incubate overnight at 4°C. Wash the slides with PBS for 3 minutes × 3 times, remove PBS, apply biotinylated goat anti-mouse/rabbit IgG polymer, and incubate at room temperature for 10 minutes. Wash the slides with PBS for 3 minutes × 3 times, remove PBS, apply streptavidin-horseradish peroxidase, and incubate at room temperature for 10 minutes. Apply freshly prepared DAB staining solution and incubate at room temperature for 2 minutes.

3.3.5 Mounting:

Slides are counterstained with hematoxylin, incubated at room temperature for 3 minutes, followed by decolorization in hydrochloric acid ethanol for 10 seconds, and then rinsed under running water for 15 minutes. Slides are dehydrated using a reverse deparaffinization process: 75% ethanol for 2 minutes, 95% ethanol for 2 minutes, absolute ethanol for 2 minutes, and finally xylene substitute for 5 minutes, with each step repeated twice. After thorough ventilation drying, slides are mounted with neutral resin.

3.3.6 Immunohistochemistry Scoring:

The scoring is based on the staining extent of positive tumor cells in the tissue: <25% scores 1 point, 25%~50% scores 2 points, 50%~75% scores 3 points, and ≥75% scores 4 points. Staining intensity is scored as follows: no staining (0 points), weak positive (1 point), moderate positive (2 points), and strong positive (3 points). The final score is obtained by multiplying the scores of staining extent and intensity. High expression is defined as 5 points or above, while low expression is less than 5 points.

3.3 **Real Time PCR**
    3.3.1 RNA Sample Preparation:

Total RNA extraction was conducted using a cell total RNA extraction kit. Specifically, 500 μL of Buffer RL was added to each well of a 6-well plate to lyse cells. After thorough cell lysis, the lysate was transferred to gDNA filter columns and centrifuged at 12000 rpm for 30 seconds to collect the filtrate. Subsequently, 250 μL of ethanol was added, mixed well, and transferred to RNA adsorption columns, followed by centrifugation at 12000 rpm for 30 seconds, and the flow-through was discarded. Washing steps were performed with 700 μL of Buffer RW1 and Buffer RW2, each followed by centrifugation and discarding of the flow-through. Finally, 500 μL of Buffer RW2 was added, centrifuged at 12000 rpm for 2 minutes, and the flow-through was discarded. The RNA adsorption column was transferred to a 1.5 mL centrifuge tube, and 50 μL of RNase-free ddH2O preheated to 65°C was added to the center of the column. After incubating at room temperature for 2 minutes, RNA was eluted by centrifugation at 12000 rpm for 1 minute, and the concentration was measured.

3.3.2 RNA Reverse Transcription:

Each sample tube containing 2 μg of RNA template was adjusted to a volume of 7 μL with DEPC water. The mixture was denatured at 65°C for 5 minutes and then chilled on ice for 2 minutes. Reverse transcription reactions were performed using the HiFi-MMLV cDNA first-strand synthesis kit. Each reaction was set up with 4 μL of dNTP Mix, 4 μL of 5×RT Buffer, 2 μL of Primer Mix, 2 μL of DTT, and 1 μL of HiFi-MMLV. The thoroughly mixed 13 μL reaction mixture was added to 7 μL of RNA template, followed by incubation at 42°C for 50 minutes and then at 85°C for 5 minutes to complete reverse transcription.

3.3.3 Real-time Fluorescent Quantitative PCR (RT-PCR):

The obtained cDNA was diluted 2-fold for use as RT-PCR templates. Each reaction well was filled with 10 μL of 2×UltraSYBR Mixture, 0.6 μL each of upstream and downstream primers for the target gene, and 6.8 μL of ddH2O. After thorough mixing and centrifugation, 2 μL of diluted cDNA template was added to each reaction well, followed by the addition of 18 μL of the above mixed system. The RT-qPCR program consisted of three steps: initial denaturation at 95°C for 10 minutes, followed by 40 cycles of PCR amplification (95°C for 10 seconds, 60°C for 30 seconds, and 72°C for 32 seconds), and finally, a melting curve analysis (95°C for 15 seconds, 60°C for 60 seconds, 95°C for 15 seconds, and 60°C for 15 seconds).

3.4 **RNA-binding protein immunoprecipitation**
    3.4.1 Sample Preparation:

MGC803 cells transfected with pcDNA3.1-ZFP36L1-Flag plasmid were washed with pre-chilled PBS at 4°C, harvested, and transferred to 1.5 mL EP tubes. After centrifugation at 1500 rpm for 5 minutes at 4°C, the supernatant was discarded. Each tube was resuspended in 250 μL of RIP lysis buffer, homogenized, and left on ice for 5 minutes. The collected cell lysates were stored at -80°C.

3.4.2 Magnetic Bead and Antibody Binding:

Fifty microliters of resuspended magnetic beads were added to each 1.5 mL EP tube, followed by the addition of 500 μL of RIP wash buffer. The tubes were vortexed and placed on a magnetic separator until the solution cleared, and the supernatant was discarded. The beads were washed again with RIP wash buffer. After resuspending the beads in 100 μL of RIP wash buffer, approximately 8 μg of Flag antibody or IgG was added to each tube, and incubated at room temperature for 30 minutes. The tubes were placed on a magnetic separator until the solution cleared, and the supernatant was discarded. The beads were washed again with RIP wash buffer.

3.4.3 RNA-binding protein immunoprecipitation:

Each tube was supplemented with 900 μL of RIP immunoprecipitation buffer. The cell lysates prepared in the first step were thawed, centrifuged at 14000 rpm for 10 minutes at 4°C, and a portion of the sample was retained as a 5% input control (Input) for direct RNA purification. One hundred microliters of the supernatant was added to the magnetic bead-antibody complex to make a total volume of 1 mL, and incubated overnight at 4°C. The tubes were placed on a magnetic separator, the supernatant was discarded, and the beads were washed six times with RIP wash buffer.

3.4.4 RNA Purification:

The immunoprecipitation complexes were resuspended in 150 μL of Proteinase K buffer and incubated at 55°C for 30 minutes. After placing the tubes on a magnetic separator, the supernatant was collected, and 250 μL of RIP wash buffer was added to each tube. Four hundred microliters of phenol-chloroform-isoamyl alcohol mixture was added, vortexed for 15 seconds, and centrifuged at 14000 rpm for 10 minutes at room temperature. The upper aqueous phase (350 μL) was carefully transferred to new EP tubes, and each tube was supplemented with 50 μL of Salt Solution I, 15 μL of Salt Solution II, 5 μL of Precipitate Enhancer, and 850 μL of ethanol. After incubating overnight at -80°C, the tubes were centrifuged at 14000 rpm for 30 minutes at 4°C, and the supernatant was discarded. The RNA pellet was washed with 80% ethanol, air-dried, and dissolved in 20 μL of DEPC-treated water.

3.4.5 RT-PCR Quantitative Analysis:

RNA was reverse transcribed into cDNA, and real-time fluorescent quantitative PCR was performed to detect the products. After normalizing the results with Input, fold differences were analyzed.

3.5 **Co-immunoprecipitation**
    3.5.1 DYKDDDDK-G1 Affinity Resin:

Resin Pre-treatment: Take 50 μL of affinity resin slurry and place it in a 1.5 mL EP tube. Add 500 μL of TBS buffer, centrifuge at 6000 g for 30 seconds, discard the supernatant, and repeat the wash step twice. Resin-Sample Binding:Add 400 μL of prepared protein sample to the resin, mix thoroughly, and incubate overnight at 4°C on a rotary mixer. Add 500 μL of TBS buffer, centrifuge at 6000 g for 30 seconds, discard the supernatant, and repeat the wash step twice. Denaturing Elution:Add 25 μL of Loading Buffer, mix well, heat at 100°C for 5 minutes, centrifuge at 6000 g for 30 seconds, collect the supernatant, and proceed with Western Blot detection.

3.5.2 Protein A/G Magnetic Bead Method:

Bead/Antibody Pre-treatment: Take 30 μL of magnetic beads and place them in a 1.5 mL EP tube. Add 400 μL of binding/washing buffer, mix well, magnetically separate, discard the supernatant, and repeat the wash step twice. Dilute the antibody with binding/washing buffer to a final concentration of 5 μg/mL. Antibody-Bead Binding: Add the diluted 400 μL antibody to the prepared beads, mix well, and incubate at 4°C for 2 hours on a rotary mixer. Magnetically separate, collect the beads, add 400 μL of binding/washing buffer, mix well, magnetically separate, discard the supernatant, and repeat the wash step twice. Antigen-Antibody-Bead Complex Binding: Add 400 μL of prepared antigen sample to the beads, mix well, and incubate at 4°C for 2 hours on a rotary mixer. Magnetically separate, collect the beads, add 400 μL of binding/washing buffer, and repeat the wash step twice. Denaturing Elution: Separate the beads, discard the supernatant, add 25 μL of Loading Buffer, mix well, heat at 95°C for 5 minutes, separate the beads, collect the supernatant, and proceed with Western Blot detection.

3.6 **GST pull-down**
    3.6.1 Protein Prokaryotic Expression:

pET-32a-His-BRD4 plasmid, pGEX-4T-2-GST-SPI1 plasmid, and empty vector plasmid were transformed into competent cells. During the logarithmic growth phase of the cultured bacteria after 5 hours of shaking, isopropyl β-D-1-thiogalactopyranoside (IPTG) was added to a final concentration of 1 mmol/L to induce expression at 18°C for 24 hours. Additionally, 500 μL of bacterial culture was taken without induction and mixed with loading buffer for subsequent experimental controls. Bacterial pellets were collected by centrifugation, resuspended in PBS containing protease inhibitors, treated with lysozyme to a final concentration of 2 mg/mL at 4°C for 30 minutes, followed by sonication at 25% power for 1 minute (3 seconds on, 3 seconds off, repeated 10 times) until clear. Centrifugation was performed at 15000 rpm for 15 minutes at 4°C.

3.6.2 His-BRD4 Protein Purification:

The cleared supernatant after centrifugation was added to Ni-NTA purification columns pre-equilibrated with PBS at a 10-fold column volume, and allowed to flow through naturally for 8 times. Gradient washing was performed using PBS containing 20 nM and 40 nM imidazole, followed by protein elution using PBS containing 250 nM imidazole. The purified protein was collected and stored at -80°C.

3.6.3 GST-SPI1 Protein Purification:

The cleared supernatant after centrifugation was incubated with GST beads for 1 hour, followed by centrifugation at 2000 g for 3 minutes, and discarding the supernatant. The GST beads were washed three times with 10-fold volume PBS, followed by elution with 1 mL GST Elution Buffer for 10 minutes, and centrifugation at 2000 g for 3 minutes. The eluate was collected, and the purification was repeated twice. The purified protein was stored at -80°C.

3.6.4 GST Pull-down Assay:

The GST pull-down assay was conducted using a GST pull-down kit, following the instructions provided. This included equilibration of glutathione resin, binding of the target protein expressed as GST fusion, preparation of prey protein, elution of bait-prey protein complex, and gel electrophoresis analysis.

3.7 **RNA pull-down**
    3.7.1 Plasmid Construction and Transfection:

pcDNA3.1-ZFP36L1-Flag and pcDNA3.1-mutZFP36L1-Flag plasmids were transfected into MGC803 cells. After 48 hours, cells were harvested and lysed using lysis buffer.

3.7.2 RNA Probe Preparation:

The clone plasmid containing the HDAC3 probe sequence was transcribed into single-stranded RNA probes using T7 RNA polymerase. After phenol-chloroform purification, the RNA probes were biotinylated using the RNA 3’ End Biotinylation Kit and incubated overnight at 16°C. RNA was then recovered using phenol-chloroform extraction and dissolved in DEPC-treated water.

3.7.3 RNA-Protein Pull-down Assay:

This assay includes pre-treatment of magnetic beads, binding of biotinylated RNA probes to streptavidin-coated magnetic beads, RNA-protein binding, washing and elution of the RNA-protein complex, positive probe controls, and Western blot analysis of the results.
